# Detail: species-agnostic, generalizable animal behavior analysis using foundation model embeddings

**DOI:** 10.64898/2026.11.24.769032

**Authors:** Moses Lurbur, Vikram Iyer, René Just, Kevin R. Coffey, Sam A. Golden

## Abstract

Automated tools for quantifying and classifying animal behavior from video data are crucial for neuroscience, medicine, and biology. Automated analysis of animal behavior from video usually starts with pose estimation, which needs manually annotated training frames for each new species or setup and reduces an animal to a small number of biased keypoints. Here we present Detail, a behavior analysis package built on foundation vision models, for species- and context-agnostic analysis of animal behavior. Animals are segmented from video frames using SAM 3, egocentrically aligned using keypoints placed by DINOv3 patch matching, and encoded by a frozen DINOv3 model as high-dimensional pose embeddings. Beyond a few annotated example frames, no labeled training data are required. Segmentation and keypoint models can be distilled into YOLO models that run quickly on a single consumer graphics card. Analysis modules operate directly on the pose embeddings for unsupervised behavior discovery, example-based behavior retrieval, precision auditing, and statistical comparison across experimental conditions. At the core of Detail is a pose embedding space, generated from an experiment’s videos, that represents animal behavior. The complete workflow, from installation to analysis, is available through graphical and command-line interfaces for scripted and high-throughput use. Detail is available at github.com/detail-embedding/detail.

## Introduction

Quantifying animal behavior is central to neuroscience, medicine, and biology. Computer vision tools now automate much of the detection, tracking, and analysis of behavior from video [1–3]. The predominant approach is pose estimation, the localization of predefined body parts (keypoints) based on manually annotated training data [4–6]. Tools such as DeepLabCut [1], SLEAP [2], and Lightning Pose [3] are widely used because they provide accurate, interpretable kinematic measurements. The resulting key-points support supervised behavior classification with tools such as SimBA [7] and unsupervised behavior discovery with B-SOiD [8], VAME [9], and Keypoint-MoSeq [10].

Keypoint-based analysis has two limitations. First, accurate keypoint detection for a new setup or species often requires manually labeled frames for fine-tuning, despite general-purpose models such as SuperAnimal [11]. Second, the selection of keypoints is both experimenter-biased and sparse. Models detect a small set of predefined points, which limits how much of an animal’s posture and behavior they can capture. Those points are chosen in advance by the user, who must judge that they capture both anticipated and unanticipated behaviors.

Foundation vision models offer an alternative. Models such as Segment Anything Model 3 (SAM 3) [12] and DINOv3 [13] have hundreds of millions to billions of parameters and are trained on internet-scale datasets (1.7 billion images for DINOv3). They learn image representations that transfer across diverse im-agery and support segmentation, object recognition, and species classification.

These models are increasingly applied to behavior analysis (Table 1). Vision transformers (ViTs) [14] have been pretrained on experimental video for pose estimation, action segmentation, and neural encoding [15]. Frozen video models have been used for behavior classification, retrieval, and localization across species [16]. Models of the SAM family [12, 17, 18] are used to detect, track, and represent animals, including from text prompts [19], and SAM 3 labels have been distilled into lightweight YOLO detectors [20].

**Table 1.** Related tools. Tools for automated analysis of animal behavior from video, from pose-based discovery to foundation-model approaches, grouped by approach, with Detail last. Preprint indicated by ^*†*^; not applicable to method indicated by —.

| Method | Species-specific training | Isolating the animal | Representation | Unsupervised discovery |
|---|---|---|---|---|
| Selfee [31] | Self-supervised (unlabeled video) | None (whole frame) | Self-supervised CNN trained on the target videos | Clustering; anomaly detection |
| Sun et al.<sup>†</sup> [16] | None | None (whole frame) | Frozen video model (VideoPrism), compared with an image model (CLIP) | — |
| BEAST [15] | Self-supervised (unlabeled video) | None (whole frame) | Vision transformer pretrained on each experiment's video | — |
| SIPEC [32] | Supervised (labeled frames) | Mask R-CNN trained on the target species; identity network | CNNs trained with behavior labels on raw frames | — |
| Annolid<sup>†</sup> [19] | Supervised (labeled frames) | Grounding DINO and SAM with text prompts; Cutie tracking | Mask R-CNN trained on labeled behavior instances | — |
| CASTLE<sup>†</sup> [21] | None (point clicks) | SAM with point prompts; DeAOT tracking | Frozen DINOv2; patch features pooled within the mask | UMAP and DBSCAN on short windows; interactive refinement |
| SingleBehavior Lab<sup>†</sup> [23] | None (point clicks) | SAM 2.1 with point prompts; SAM 2.1 tracking | Frozen VideoPrism on registered, background-removed clips | Leiden or HDBSCAN on 13-frame clips |
| MouseGPT<sup>†</sup> [25] | Supervised (labeled frames) | None (whole frame) | Mouse-specific VLM trained on more than 42 million frames | Agent-driven clustering |
| BehaveAgent<sup>†</sup> [26] | None | Molmo points from text prompts; SAM 2 tracking | Mask shape and optical-flow features over 0.4 s | UMAP clusters labeled by a VLM; LLM/VLM temporal segmentation |
| BehaviorVLM<sup>†</sup> [27] | Unsupervised (unlabeled video) | None (whole frame) | Fused video and keypoint features (LookAgain) | Deep embedded clustering, then LLM merging |
| Keypoint-MoSeq [10] | Supervised (labeled frames) | Pose estimation (keypoints) | Keypoint trajectories | Autoregressive HMM syllables |
| Faria et al.<sup>†</sup> [20] | Automatic (SAM 3 labels) | SAM 3 labels distilled into YOLOv8 | — | — |
| **Detail (this work)** | None (text prompt; optional keypoint clicks) | SAM 3 with text prompts, optionally distilled into YOLO11; mask tracking | Frozen DINOv3 embedding of the masked, egocentrically aligned animal, per frame | Prototype codebook; HMM/Viterbi bouts; average-linkage tree |

Several tools combine segmentation with a vision embedding model to isolate the animal and represent its posture. CASTLE [21] uses SAM to segment animals or body parts marked by the user and clusters DINOv2 [22] features pooled within the mask. SingleBehavior Lab [23] segments animals with SAM 2 and encodes clips of the isolated animal with the video model VideoPrism [24] for unsupervised clustering and downstream classification. A separate line of work applies vision–language models (VLMs) to behavior annotation, either by fine-tuning general-purpose VLMs or by using existing open or commercial models, with little task-specific training [25–27].

Here we present Detail, a software package for species-agnostic animal behavior analysis. Like CASTLE and SingleBehavior Lab, Detail isolates animals from experimental footage and represents them with a vision embedding model. It improves on currently available platforms in four respects: (i) *Text-prompted segmentation*. Animals are segmented from a text prompt with SAM 3, so no point prompts or manual labels are required. (ii) *Aligned pose embeddings*. Each segmented animal is egocentrically aligned before embedding, using keypoints that DINOv3 patch matching places from a few annotated frames. The result is a pose embedding that is largely invariant to the animal’s position and orientation: a 1,024-dimensional representation in place of a small set of keypoints. (iii)*Analysis on the embeddings*. Analysis modules operate directly on the pose embeddings for unsupervised temporal clustering, example-based behavior querying, precision auditing, and statistical comparison across experimental conditions. (iv) *Distillation*. Segmentation and keypoint models can be distilled [20, 28, 29] into YOLO models [30] that run faster than real time on consumer graphics processing units (GPUs). Critically, Detail brings all these components together in a single interface that guides users of any experience level from setup to analysis.

The complete workflow, from installation and model management through processing, analysis, and export, is available through a graphical interface, and through a command-line interface for scripted and high-throughput use. Detections, tracks, keypoints, pose embeddings, behavior labels, and figures can be exported in standard formats for external analysis.

We illustrate the workflow on top-down recordings of rats given different doses of fentanyl and naloxone precipitated withdrawal. Planned extensions include multi-animal interactions, video collected in natural environments, and further analyses of the pose embedding space.

## Results

### Using Detail

Detail enables users to go from experimental videos to behavior labels in a single tool without experiment-specific annotation or training (Figure 1). Detail is designed to be easy to use for those with no programming or machine learning experience, while also allowing users precise control over advanced parameters in the graphical user interface (GUI) and command-line interface (CLI). That said, we recognize that users of behavioral analysis platforms come from interdisciplinary backgrounds with varying levels of comfort with using open-source packages; we encourage users to reach out by posting an Issue on the Detail GitHub if they have questions or improvements for the installation or use of Detail.

**Figure 1.**
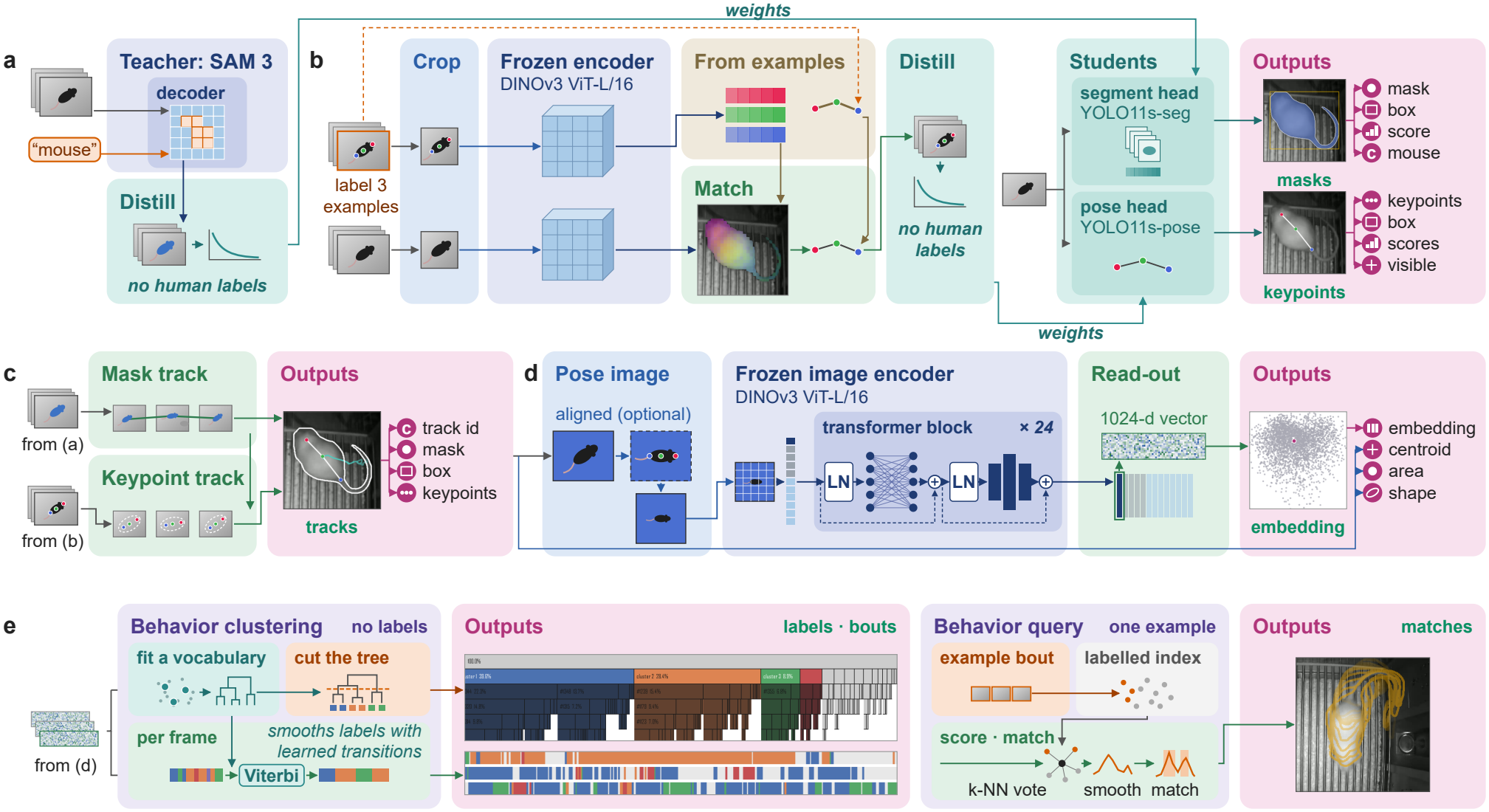
Detail for behavioral analysis. Each row runs from inputs (left) to outputs (pink). Pane color marks the kind of step: orange, user input; light blue, a fixed image operation; dark blue, a frozen pretrained network; sand, statistics of points the user clicked; teal, fit to the data without human labels; green, a fixed rule. Outputs are real results from a demonstration recording. **a**, Masks. With a text prompt (“mouse”), the frozen SAM 3 teacher segments a sample of varied frames (1,000 by default) drawn across experimental recordings. These masks train a YOLO11s-seg student, which segments every experimental frame (mask, box, score and class). **b**, Keypoints. The user clicks *n* points on up to three example frames. Crops of the examples and of sampled frames are encoded by frozen DINOv3 ViT-L/16. Features at the clicked points form a feature bank, and distances between points form edge priors. Each sampled frame’s patch features are compared with the bank by cosine similarity, giving one map per point that is decoded into a skeleton under the priors. The panel shows a sampled frame’s patch features, colored by their first three principal components and overlaid on the animal. These keypoints train a YOLO11s-pose student, run on every experimental frame. **c**, Tracking. Detections are linked into a single-subject track (mask track). Each keypoint is assigned to the tracked animal by its mask and smoothed over time (keypoint track). **d**, Embedding. The tracked animal is cropped, masked onto a uniform background, optionally rotated to its heading from **b** based on keypoints, and centered at 224 *×* 224 pixels. The class token of the frozen DINOv3 encoder gives a 1,024-dimensional embedding per frame (output: a two-dimensional PCA of one recording). Centroid, area and shape are measured from the mask. **e**, Left, clustering without labels. A vocabulary of codewords is fit to temporal windows of the embeddings and arranged as a tree, which the user cuts to set the number of behaviors. Each frame takes its nearest codeword, and a Viterbi decode with transition probabilities fit to the data smooths the sequence (outputs: the cluster tree and the ethogram of three recordings). Right, query from one example. Frames of an example bout and 500 background frames form a labeled index. Each frame is scored by a soft *k*-nearest-neighbor vote; the score is smoothed, and each match extends from its peak to half the peak score (output: a match to a rearing query, with mask outline and 0.5-s trail).

#### Documentation

Complete documentation for Detail can be found on our documentation page (detail-embedding.github.io/detail). In addition to installation and user guides, we also provide a complete application programming interface (API) reference to support external contributions and the use of Detail with agentic tools.

#### Installation

Installation, and assessment of hardware requirements, is straightforward and detailed in the User Guide. Briefly, after cloning the repository (github.com/detail-embedding/detail) and creating a Python virtual environment using uv, users are guided by the GUI through GPU configuration, Hugging Face integration, and model installation. Detail automatically diagnoses CUDA and PyTorch configuration issues and generates terminal commands to resolve them. Detail also walks users through other common difficulties in machine learning workflows, such as installing SAM 3 from source code, setting up Hugging Face model access, and installing model weights.

#### GUI and CLI

Detail can be used through a GUI or a CLI. The GUI is designed to be approachable and easy to use (Figure 2). The CLI gives advanced users granular control over every step of the data processing and analysis pipeline and makes the tool usable on headless virtual machines or with agentic tools.

**Figure 2.**
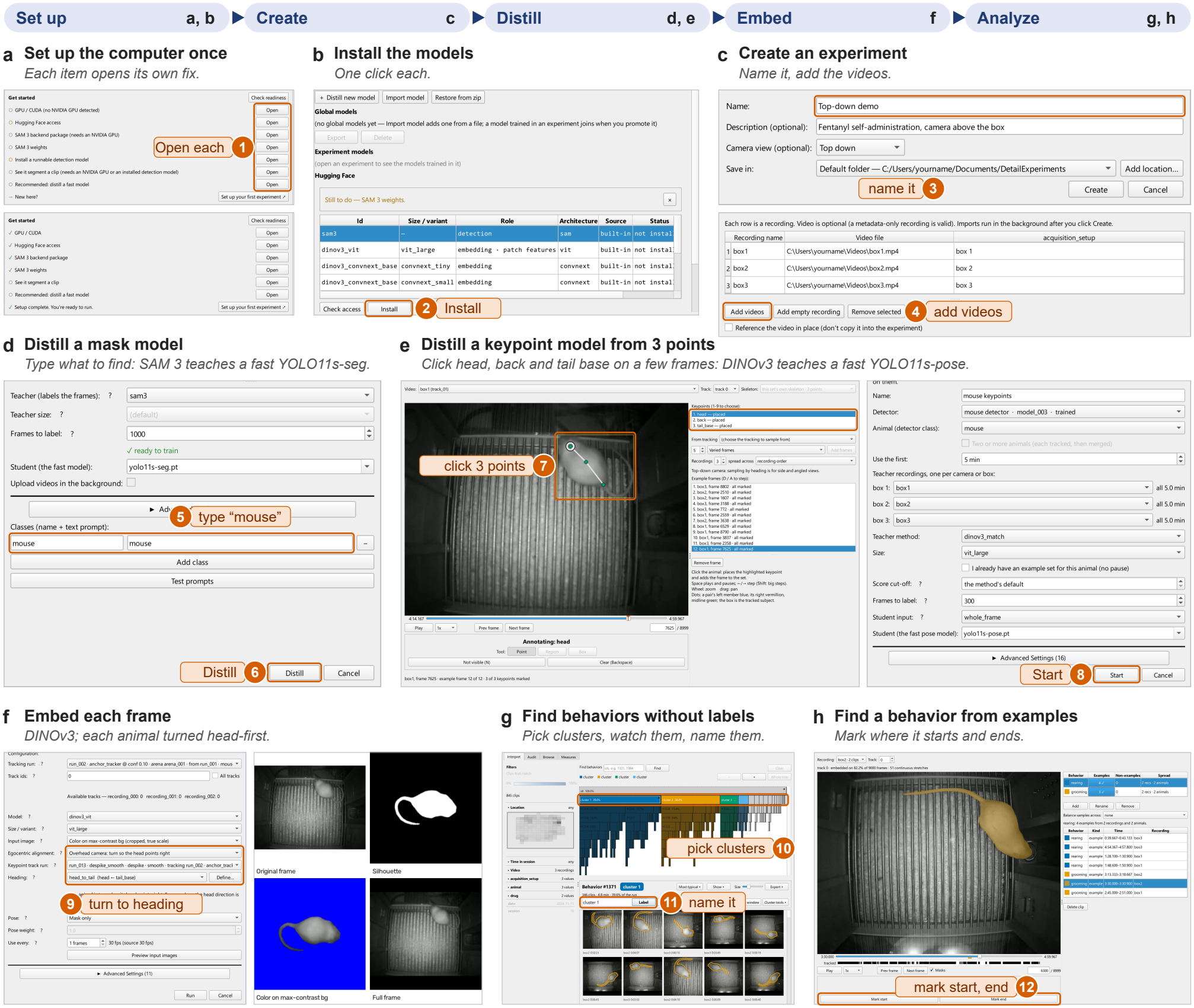
Using Detail’s graphical interface. Cropped screens of the Detail package, in the order a user meets them, from a demonstration experiment. Orange outlines and numbered callouts mark some things the user does; the package automates most else. **a**, The setup checklist before (top) and after (bottom) a computer is configured; each item’s Open button leads to the page that resolves it. **b**, The model manager; built-in models install with one click. **c**, Creating an experiment: a name, then the videos. **d**, Distilling a mask model: the user types what to find (“mouse”); the frozen SAM 3 teacher labels sampled frames, and a fast YOLO11s-seg student is trained on them. **e**, Distilling a keypoint model: the user clicks example keypoints (head, back and tail base here) on a few example frames (left); the frozen DINOv3 patch-matching teacher labels sampled frames, and a fast YOLO11s-pose student is trained on them (right). **f**, Embedding: frozen DINOv3 embeds every frame, each animal turned head-first for egocentric alignment using the heading from its keypoint track (left); right, one frame as four of the input-image options render it (original frame, silhouette, color on a maximum-contrast background, full frame). **g**, Finding behaviors without labels: the user picks clusters on the cluster tree, reviews their clips and names them. **h**, Finding a behavior from examples: the user marks where example bouts start and end.

#### Metadata, stream data, and scene geometry integration

Detail enables the integration and use of a variety of experimental data types in addition to videos. Supported data include experimental metadata (e.g. animal ID, treatment) and stream data (e.g. operant events, sensor data, manual behavior annotations).

Experimental metadata can be manually entered in Detail or bulk imported, and used to filter the results of analyses such as temporal clustering. Stream data, including any comma-separated values (CSV) time series (like physiological or neural recordings) or event data (like manual annotations), as well as supported formats like Med Associates MED-PC and BORIS [33], can be aligned to video timestamps at import. Both stream data and experimental metadata can be exported alongside processed outputs, such as detection results, for downstream analysis. Analysis modules that directly integrate stream data with behavior clusters are in development.

Users can also define detailed scene geometry in their videos to improve tracking results, provide descriptive statistics of tracked animals (location, distance, velocity, etc.), and support external data analysis. Within the GUI, users can define a field of view to restrict tracking to an area of the video. Users can also define regions or points of interest and distance calibrations, as well as batch-copy all settings to all recordings within an experiment. Users can then export the location and calibrated distance of subjects relative to the regions and points. A dedicated Location Export analysis module is provided for in-depth analysis and visualization of locomotor metrics based on scene geometry. Analysis modules that directly integrate points and regions of interest with behavior clusters are in development.

### Data processing pipeline

Detail’s data processing pipeline transforms videos of animal behavior into rich mathematical representations called pose embeddings that capture detailed information about pose and behavior. The pipeline generates bounding boxes, segmentation masks, keypoint coordinates, and movement tracks that can be analyzed directly within Detail or exported for external analysis (Figure 1).

Detail makes state-of-the-art foundation models (such as SAM 3 and DINOv3) accessible for zero-shot segmentation and keypoint tracking. Users can isolate animals and locate body parts without manually labeling training data for each experiment. Because foundation models are computationally intensive, Detail features a streamlined model distillation process for object and keypoint detection that uses a small number of frames labeled by a larger foundation model to train faster YOLO models. This allows users to efficiently harness the power of foundation models on consumer hardware. In our benchmarking, a distilled YOLO11s segmentation model gave an 18-fold speedup over SAM 3, performing segmentation at 197 frames per second (fps) on an NVIDIA RTX 5090 GPU (Table 2), and agreed with SAM 3 at a mask mean average precision of 0.93 at 50% overlap (mAP50) on held-out frames. A distilled YOLO11s-pose model agreed with patch matching at a pose mAP50 of 0.98 (Table 3). Using distilled models, Detail fully processed a 1-h video in 0.30 h on the same GPU (Table 2). For scalability, full high-performance computing (HPC) cluster support via the Detail GUI will be available in the next update.

**Table 2.** Processing time for a 1-h video. Time to run the data processing pipeline on a single 1-h video (108,000 frames at 30 fps), in decimal hours and frames per second, using default settings. Total time is based on YOLO models for detection and keypoints. Tracking time is nominal and omitted. SAM 3 does not support Apple Silicon, and patch matching is too slow to be practical on the M3 chip.

|  | RTX 5090 |  | RTX 2080 Ti |  | Apple M3 |  |
|---|---|---|---|---|---|---|
|  | h | fps | h | fps | h | fps |
| *Object detection* |  |  |  |  |  |  |
| SAM 3 | 2.74 | 11 | 6.53 | 5 | – | – |
| YOLO11s | 0.15 | 197 | 0.18 | 164 | 0.74 | 40 |
| *Keypoint detection* |  |  |  |  |  |  |
| Patch matching | 0.20 | 136 | 0.58 | 50 | – | – |
| YOLO11s-pose | 0.10 | 312 | 0.13 | 240 | 0.58 | 52 |
| *Embedding* |  |  |  |  |  |  |
| DINOv3 ViT-L | 0.05 | 530 | 0.16 | 180 | 3.39 | 8 |
| Total, YOLO models | 0.30 |  | 0.47 |  | 4.71 |  |

**Table 3.**
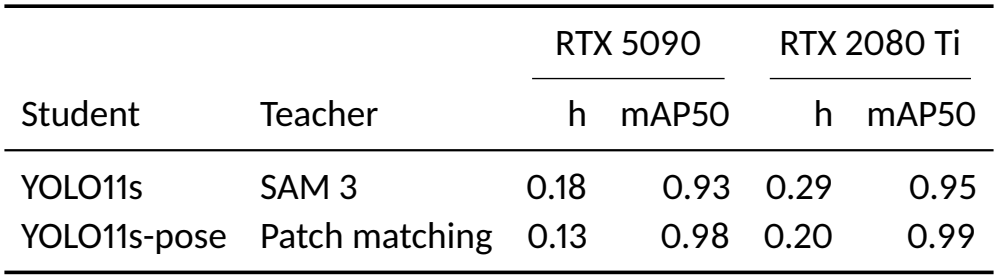
Distillation time. Time to distill the YOLO11s models used in the pipeline, in decimal hours, using default settings. Distillation is a one-time cost. Time includes labeling. Each model was trained on 1,000 frames, labeled by SAM 3 for segmentation and by patch matching for key-points. mAP50: each distilled model’s agreement with its teacher on the 15% of labeled frames held out (mask mAP50 for YOLO11s, pose mAP50 for YOLO11s-pose). Each machine trained its own model.

While Detail supports extensive species-agnostic keypoint detection and tracking, the current stated purpose of keypoints within Detail is to automatically track animal orientation (e.g. location of head and tail base), without the need to train a pose estimation model. Tracking orientation allows Detail to rotate and align the segmented animal to create an egocentric pose image prior to generating embeddings. Standardizing the animal’s orientation reduces orientation information in our pose embedding space (Table 4) and allows our downstream analysis modules to better identify changes in posture rather than orientation.

**Table 4.**
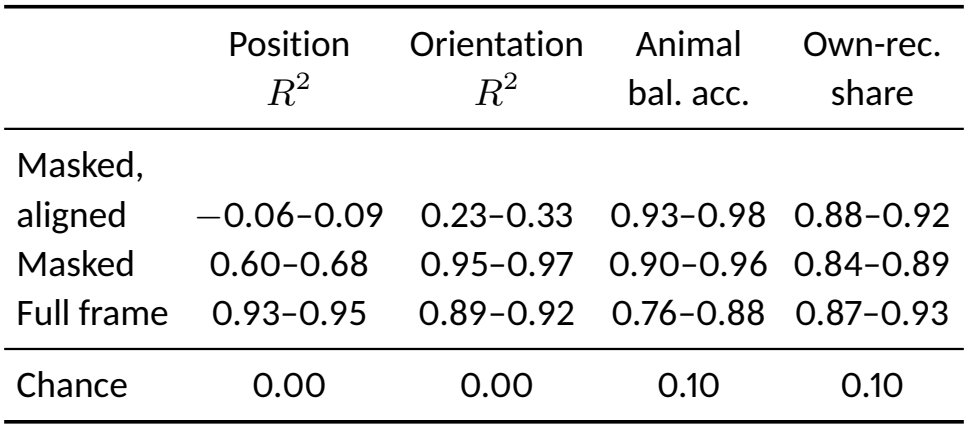
Orientation invariance of pose embeddings. Information captured in pose embeddings derived from the following pose images: masked and egocentrically aligned, masked, and full video frame. *R*^2^ values are based on MLP prediction of animal position (centroid of the mask in the video frame) and orientation (direction from the tail and midpoint keypoints to the head keypoint in the video frame). Animal balanced accuracy is the mean per-animal recall of an MLP that identifies the animal, trained on four treatments and tested on the fifth. Own-recording share is the fraction of a frame’s 10 nearest neighbors that come from its own recording. Each cell is a 95% confidence interval from 1,000 bootstrap resamples of animals. Chance is the expected score if the pose embedding carried no information for the given variable. We use 10 animals, 50 recordings, and 2,000 evenly spaced frames per recording for our estimates.

|  | Position<br>$R^2$ | Orientation<br>$R^2$ | Animal<br>bal. acc. | Own-rec.<br>share |
|---|---|---|---|---|
| Masked,<br>aligned | −0.06–0.09 | 0.23–0.33 | 0.93–0.98 | 0.88–0.92 |
| Masked | 0.60–0.68 | 0.95–0.97 | 0.90–0.96 | 0.84–0.89 |
| Full frame | 0.93–0.95 | 0.89–0.92 | 0.76–0.88 | 0.87–0.93 |
| Chance | 0.00 | 0.00 | 0.10 | 0.10 |

To generate pose embeddings from a video, the pipeline executes the following steps:

1. **Segment** the subject from the background to create a segmentation mask.
2. **Track** the subject across video frames.
3. **Detect and track keypoints** to rotate the subject’s mask into a consistent, egocentric alignment.
4. **Generate a pose embedding** from the aligned pose image that captures high-dimensional information about pose.

#### Object detection and tracking

The pipeline begins with object detection, where Detail uses a text-promptable segmentation model (SAM 3) to detect the subject of interest. SAM 3 is a foundation vision model that, given a text string such as “rat” or “fish”, produces bounding boxes and pixel-level masks of objects matching that string (Figure 3).

**Figure 3.**
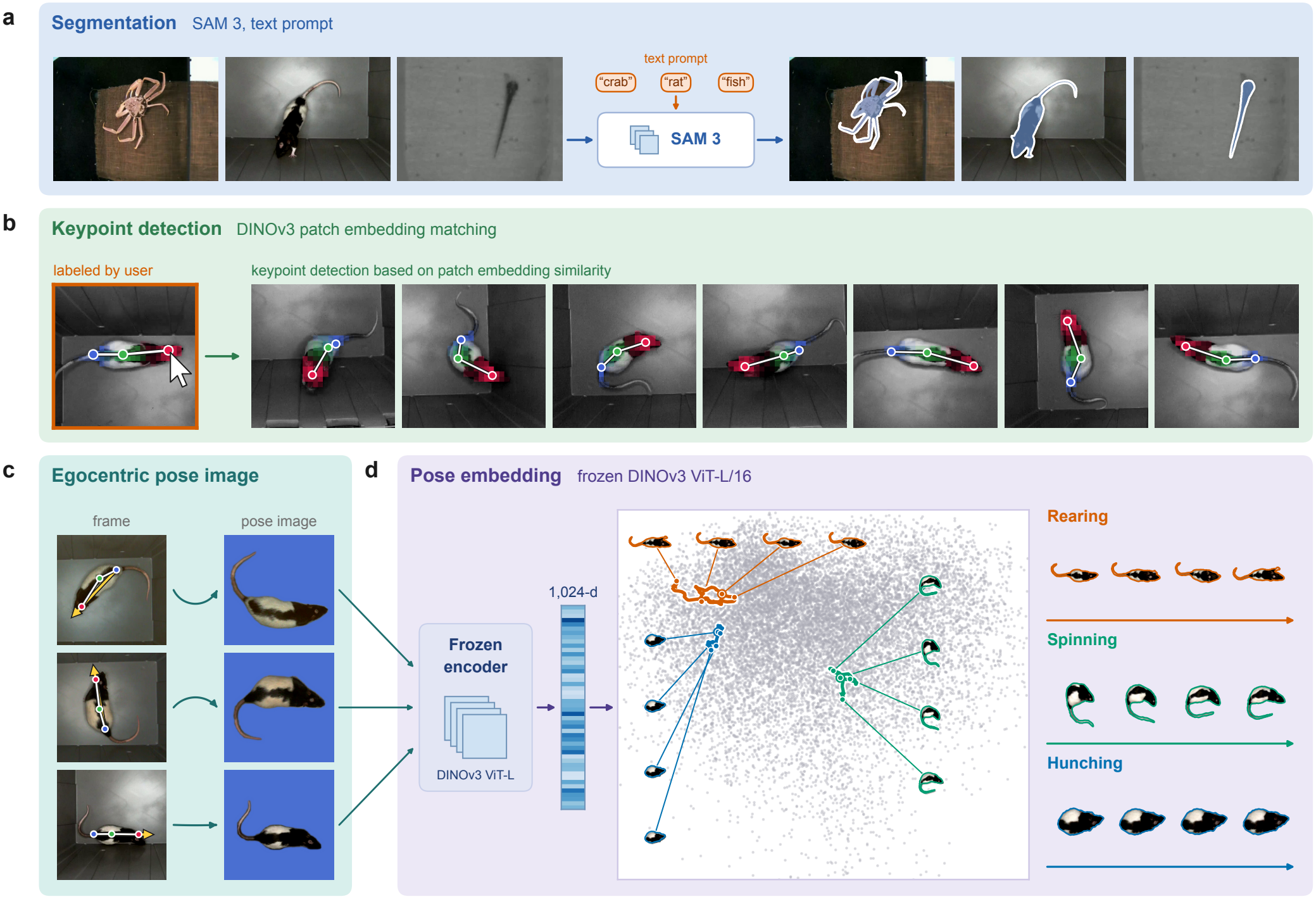
Detail’s pipeline steps from segmentation to pose embedding. **a**, Segmentation: SAM 3 segments the subject of interest based on a text prompt, the only species-specific input in our pipeline. **b**, Keypoint detection by patch matching: a user labels head, midpoint and tail on a few frames. Similarity of DINOv3 patch embeddings is used to label the rest of the experiment. Color shows each patch’s similarity to the labeled patch. **c**, The egocentric pose image is built by orienting every mask to face right on a high-contrast background. **d**, Pose embedding: DINOv3 encodes each pose image as a 1,024-d vector to build the pose embedding space. The pose embedding space is shown as a 2D PCA, with the path through it for one bout of rearing, spinning, and hunching shown.

We have found SAM 3 to be very generalizable and to produce accurate detections across a wide array of animals and experimental setups. Using multi-class prompts (e.g. “mouse” and “cable”) is supported to remove detection artifacts from complex recording scenes. After detection, we track objects across frames, which acts as an error-filtering step, removing spurious detections and unifying animal identity across frames.

#### Keypoint detection and tracking

Detail uses DINOv3-derived keypoints to egocentrically align the pose image prior to embedding. The default skeleton is three points along the midline (head, midpoint, and tail base), which are used to define the animal’s orientation. Users can edit, create, or import other skeletons, and keypoints can be exported for external use.

For species- and setting-agnostic keypoint detection, we use patch matching, which compares the similarity of patch embeddings of an image embedding model (DINOv3), to detect keypoints across different videos and animals (Figure 3). Users first annotate key-points on a small number of video frames (we suggest 1–3 example frames). For every tracked frame, Detail crops the animal, computes DINOv3 patch embeddings for the crop, and places each keypoint at the patch most similar to the marked patch. One marked frame of a single animal is often enough to place the three points across other animals, sessions, and orientations (Figure 3). Keypoint tracking then assigns detections to the tracked subject, removes outlier detections, and smooths the keypoint trajectories.

#### Embedding

To generate a pose image for every frame of a video, the subject is segmented from the frame using its tracked mask, egocentrically aligned using keypoints, centered, scaled, and overlaid on a solid background. To capture rich pose information, we pass the ego-centrically aligned images through an image embedding model (DINOv3) and save the resulting embedding, the model’s class (CLS) token, as the frame’s pose embedding.

#### Model distillation

Foundation models like SAM 3 are very accurate but computationally expensive and slow. Detail provides a simple workflow for training smaller, faster models on labels generated by larger models such as SAM 3. This technique is commonly referred to as model distillation [28, 29].

To distill an object detection model, we use SAM 3 to generate detections for the text string of interest, which are then used to train the smaller, faster YOLO model. For keypoint detection, we use the smoothed and tracked results from patch matching to train a small keypoint detection model.

Labeling 1,000 frames and training took 0.18 h for the segmentation model, labeled by SAM 3, and 0.13 h for the keypoint model, labeled by patch matching, on an RTX 5090 (Table 3).

### The pose embedding space

The pose embedding space is the high-dimensional space occupied by our pose embeddings. Every frame of a video is represented as a point in the pose embedding space (Figure 4). Our pose embeddings have several properties that make them strong alternatives to keypoints: high dimensionality, invariance to animal location, low sensitivity to orientation, and high cosine similarity for similar poses across animals.

**Figure 4.**
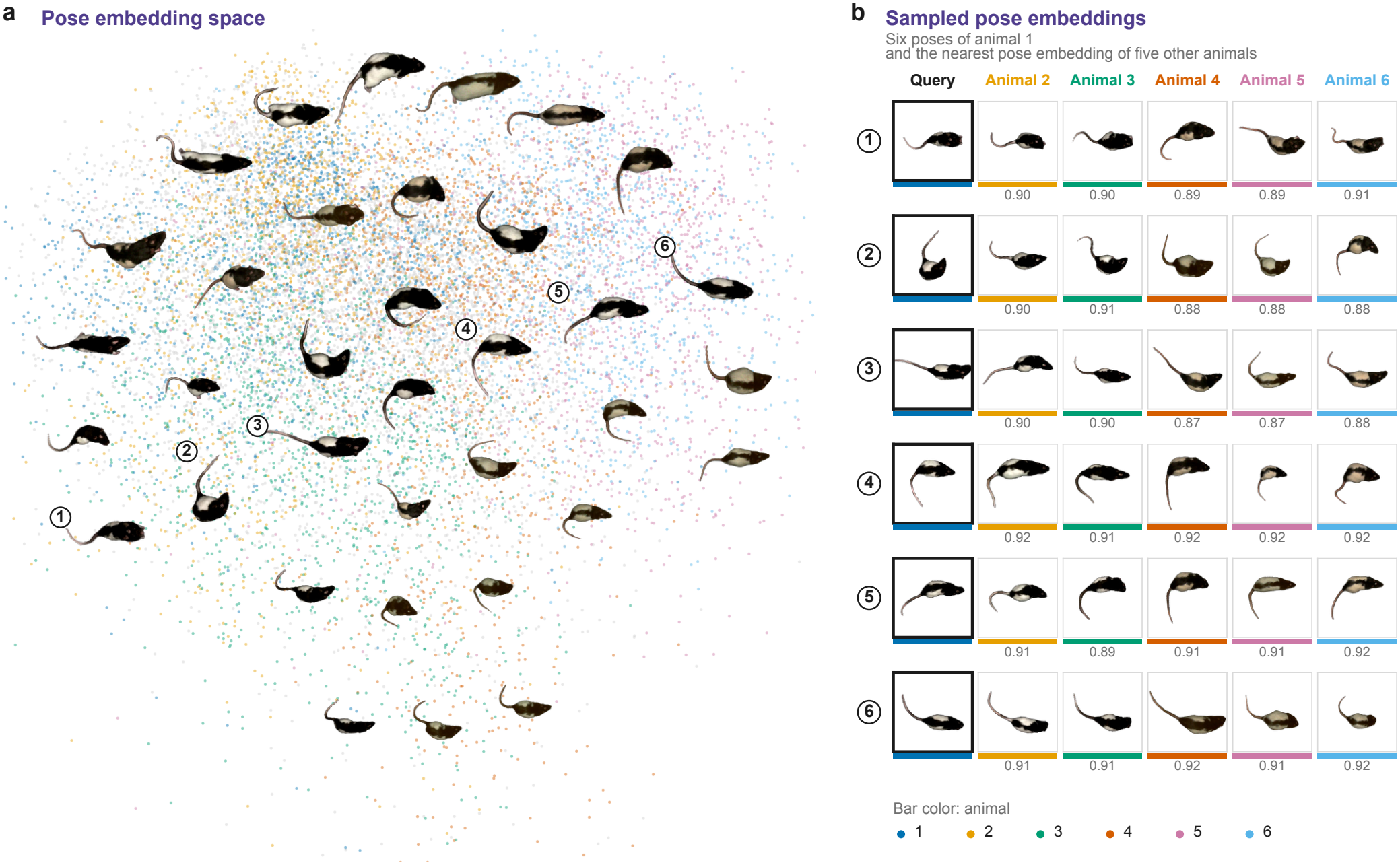
The pose embedding space visualized. **a**, The 1,024-d pose embeddings of 15,000 frames from 50 recordings of 10 rats, shown as a 2D PCA plot. The pose images of selected frames are drawn at their positions; six are numbered. **b**, The six numbered poses, all from one rat, and the nearest frame of each of five other rats by cosine similarity in the full 1,024-d space. The cosine similarity to the query is shown under each image; two frames drawn at random average 0.80 *±* 0.08. Color indicates animal ID.

Our pose embeddings have 1,024 dimensions, compared with 14 coordinates for the seven keypoints of the CalMS21 benchmark [34]. Egocentric alignment removes nearly all information about the animal’s position and orientation. A multilayer perceptron (MLP) predicts position with an *R*^2^ of *™*0.06 to 0.09 and orientation with an *R*^2^ of 0.23 to 0.33 from aligned embeddings, compared with an *R*^2^ of 0.60 to 0.68 and 0.95 to 0.97 without alignment (Table 4). Inspection of an embedding’s nearest neighbors by cosine similarity also shows that similar poses from different rats have very similar embeddings, and dissimilar poses are distant from each other (Figure 4). The embeddings do still identify the animal and source recording: an MLP tells 10 rats apart with a balanced accuracy of 0.93 to 0.98, and about 90% of a frame’s nearest neighbors come from its own recording (Table 4).

### Analysis modules

Detail has two analysis modules that use the pose embeddings produced by our data pipeline to detect, group, and discover behaviors and poses: temporal clustering and behavior querying.

#### Temporal clustering

Temporal clustering produces a hierarchy of clusters of sequential pose embeddings. The Detail GUI makes it easy to browse and label clusters, view ethograms of labeled behaviors, and filter behaviors by subject location and user-provided experimental variables like animal ID or treatment. The hierarchical structure of the clusters allows users to easily label behaviors at many levels of granularity. Future releases will focus on refining automated descriptions of behavioral clusters.

#### Behavior querying

Behavior querying allows a user to find behaviors of interest across their entire experiment. The user starts by selecting one or more examples of a behavior in their experiment. Detail computes the similarity of the example sequences with every other sequence of pose embeddings across the entire experiment, returning matches in order of similarity. Users can then inspect the results, select an appropriate confidence cutoff, and export the labeled data for downstream analysis. To aid in this process, clusters (or subsets of clusters) identified in a temporal clustering analysis can be used in a behavior query analysis, via the GUI.

#### Auditing

Accuracy metrics are an essential part of any behavioral analysis workflow. Every analysis module in Detail comes with the ability to audit the precision of classified results. The GUI allows users to quickly review every clip of a labeled behavior and remove clips that do not belong, giving an exact precision for each behavior.

### Comparisons

Comparisons allow users to compare behavior, movement, and body shape across experimental metadata, such as treatment, sex, or dose. Users can choose how recordings are grouped, and Detail selects the matching statistical design, comparing between animals (e.g. sex) or within each animal (e.g. dose). Differences are tested with permutation tests corrected for the number of behaviors compared. Users can compare labeled behaviors or every cluster from temporal clustering to find differences before labeling. Results can be exported as figures with accompanying statistics. Here, we used the comparisons module to identify behavior clusters whose prevalence changed with increasing fentanyl dose and naloxone-precipitated withdrawal (Figure 5d–f).

**Figure 5.**
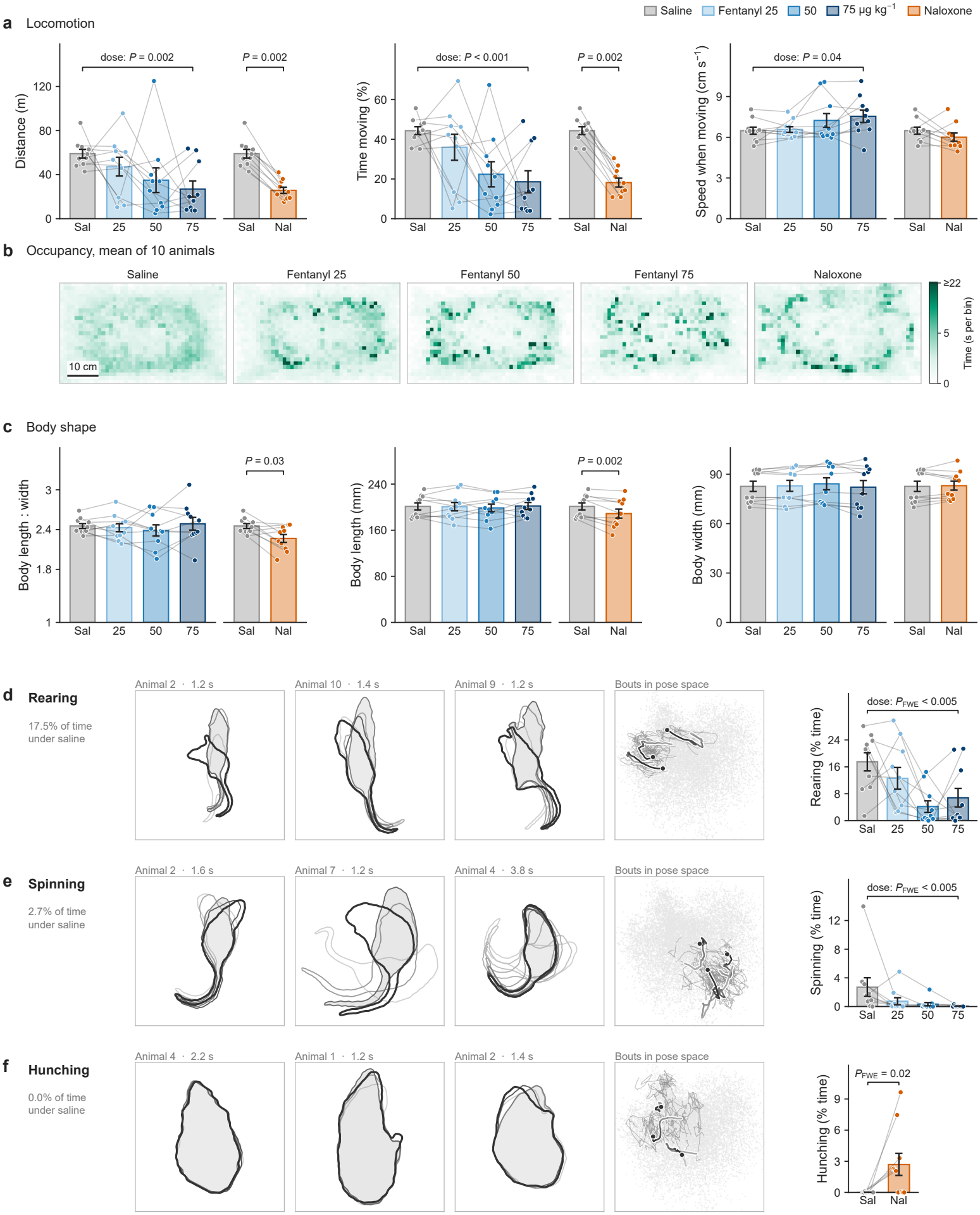
Fentanyl and naloxone change locomotion, body shape and behavior in rats recorded from above. **a**, Locomotion during a 30-min session after saline, fentanyl (25, 50 or 75 *µ*g/kg) or naloxone (1 mg/kg): distance travelled (left), share of time spent moving (middle) and mean speed while moving (right). **b**, Occupancy maps: time spent in each 1.2-cm bin, averaged over the ten rats in each session. Scale bar, 10 cm. **c**, Body shape: length-to-width ratio (left), body length (middle) and body width (right). **d–f**, Behavior clusters whose share of session time differed between conditions: rearing (**d**), spinning (**e**) and hunching (**f**). Left; three example bouts, each from a different rat. Outlines are drawn every 0.4 s from light (start) to dark (end), with the middle outline shaded, egocentrically aligned so that the head points up. Middle; the same bouts as paths through pose space, from light (start) to dark (end, dot), over the cluster’s 40 most typical bouts (thin grey lines). Pose space is the first two principal components of the features the clusters were found in (17% of their variance), the same axes in every row; grey dots are a random sample of all poses. Right; share of session time spent in the cluster. In all bar graphs, bars show mean *±* s.e.m., dots are individual rats (*n* = 10 rats, 5 female and 5 male). Brackets marked “dose” show a within-animal permutation test of the association with fentanyl dose (saline counted as 0); other brackets compare saline with naloxone by a paired permutation test. *P*_FWE_, *P* value corrected for the number of clusters screened. Only *P <* 0.05 shown. Sal=saline; Nal=naloxone.

### Data export

Detail supports comparisons within the tool, but many experiments require highly specialized analyses. Detail makes it easy to export all data produced within the tool, including segmentation masks, bounding boxes, tracks, keypoint detections, pose embeddings, and results from any analysis module. Every piece of data can be exported via the GUI or CLI.

### Experimental results

#### Fentanyl and naloxone-precipitated withdrawal produce distinct changes in movement, posture, and behavior

Fentanyl reduced locomotion in a dose-dependent manner (Figure 5a). Distance traveled fell from 59 *±* 4 m after saline to 27 *±* 7 m at 75 *µ*g/kg (*P* = 0.002), and the share of time spent moving fell from 44 *±* 2% to 19 *±* 6% (*P <* 0.001). When rats did move, they moved slightly faster at higher doses (6.5 *±* 0.3 cm/s after saline, 7.5 *±* 0.5 cm/s at 75 *µ*g/kg; *P* = 0.04). Occupancy maps help visualize these differences. After saline, rats explored the perimeter of the arena evenly, whereas fentanyl animals spent time in a few concentrated places (Figure 5b). Naloxone-precipitated withdrawal also reduced locomotion relative to saline. Distance traveled (26 *±* 3 m) and time moving (18 *±* 2%) were lower in every rat (both *P* = 0.002), with no change in speed while moving. Body shape separated fentanyl exposure and withdrawal (Figure 5c). After naloxone, rats were shorter than after saline (189 *±* 8 mm compared with 201 *±* 6 mm; *P* = 0.002) but no wider (83 *±* 3 mm in both sessions; *P* = 0.69), so their length-to-width ratio fell (2.27 *±* 0.06 compared with 2.45 *±* 0.04; *P* = 0.03). This is consistent with the compact “hunched” posture commonly associated with feelings of sickness in rats.

Temporal clustering identified the behaviors behind these changes, without our specifying them in advance (Figure 5d–f). Exploratory behavior such as rearing fell with fentanyl dose, from 17.5 *±* 2.7% of session time after saline to 6.8 *±* 2.8% at 75 *µ*g/kg, and spinning fell from 2.7 *±* 1.3% to 0.05 *±* 0.03% (family-wise *P <* 0.005). Hunching was absent after saline (0.1% of session time or less in every rat) and appeared during naloxone-precipitated withdrawal, occupying 2–10% of the session in six of the 10 rats (mean 2.7 *±* 1.1%; family-wise *P* = 0.02).

## Methods

### Software design and implementation

We designed the architecture of Detail with three goals: flexibility, reliability, and scalability.

At the core of Detail is its Python API, which we designed along with a GUI and CLI to accommodate a wide range of use cases and grow with the needs of our user base and advances in computer vision and machine learning.

Detail safely manages computationally expensive processes and allows aborted or failed jobs to be rerun without repeating completed work. We also designed the tool to ensure experimental reproducibility and good data management practices.

We have designed Detail’s architecture to make the data processing pipeline tractable to run on a wide range of hardware. The processing code has no GUI dependency, so Detail can be used headlessly via the CLI on a server or cluster. Detail also supports parallel processing on multi-GPU machines. Long-running tasks like detection and embedding are run as back-ground processes that persist when the app is closed and report live progress and estimated run time.

Every core function of Detail is designed to be modular, including data management, the data processing pipeline, and analyses. This modularity makes it simple to add new models, tracking algorithms, and analytical approaches and supports continued community development.

Detail is extensively documented. The API reference is generated from the code’s docstrings, and each stage’s parameters are defined once in a pydantic model that generates the GUI forms, the CLI help text, and the parameter reference. User-facing guides cover the GUI and CLI workflows.

#### Experiment structure

An experiment is the top-level data object in Detail. Each experiment is a self-contained project made up of recordings, experimental metadata (e.g. animal ID, treatment type) stored in one table, models trained within it, and all data generated during data processing and analysis (Figure 6). A recording is one animal in one session and holds its video, stream data (e.g. events, sensor data), and scene geometry (e.g. field of view, regions of interest, points of interest, distance calibration).

**Figure 6.**
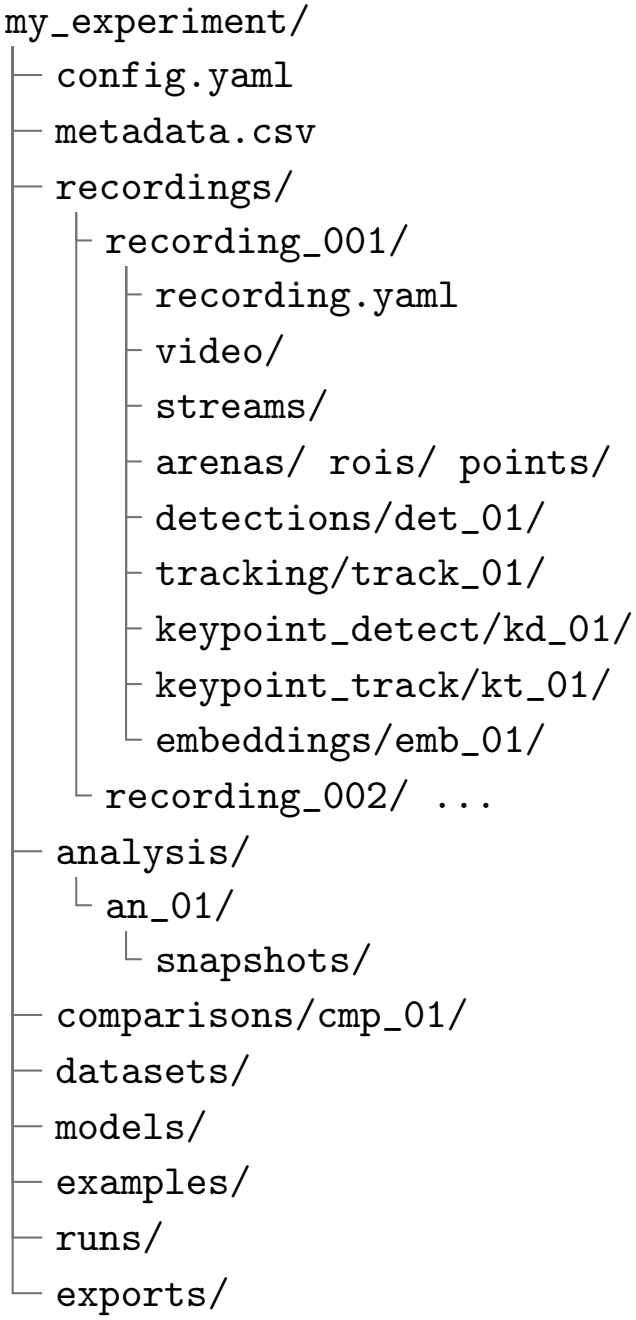
Experiment structure in Detail. Experiments are self-contained, so the folder can be copied or moved to another machine. Each recording holds its own data processing runs. Analyses and comparisons span recordings and are stored at the experiment level. Videos can be referenced in place rather than copied into the experiment.

Experiments are designed for reproducibility. Each data processing step (e.g. detection, tracking, embedding) produces a run. Every run is immutable and saves the full settings it ran with, including all parameters, model identifiers, and the upstream run it read from.

#### Data management

Detail’s management of our core data components— videos, model weights, pipeline data, and analysis results—is designed to be space-efficient and portable, and to ensure experimental reproducibility.

Pipeline data and analysis results are stored as Zarr arrays [35], and records and settings are stored as human-readable YAML files. All paths within an experiment are relative, so an experiment can be copied or moved to another machine.

Videos can be referenced in place and do not need to be copied into the experiment. Referenced video files are fingerprinted, and moved or altered videos are automatically detected and refused during data processing.

Model weights are only downloaded when explicitly requested by a user, saving space and ensuring that Detail can run offline. Trained models can be exported and imported for use by collaborators or on another machine running Detail.

Runs cannot be deleted while another step of the processing pipeline depends on their data. To ensure that results from analyses are reproducible and not accidentally overwritten, snapshots are automatically taken when data are used in an export or comparison.

#### Metadata, stream data, and scene geometry

Experimental metadata is stored in a single table with one row per recording and any number of user-defined columns. Metadata can be entered in the GUI or CLI, or bulk imported from a CSV file.

Stream data can be imported from CSV time series and event files, frame-indexed behavior annotations, MED-PC output, and BORIS annotations [33]. Streams are converted to int64 nanosecond times-tamps, and streams timed in video frames or seconds are converted to video timestamps. Streams with their own clock are aligned to videos with a constant offset, which users can set by marking the video frame where a stream event occurs.

The field of view, regions of interest, and points of interest are defined in normalized image coordinates and can be adjusted across a video using keyframes. During tracking, detections whose bounding box center is outside the field of view are excluded. Distances are calibrated based on a user-defined distance between two points. Scene geometry can be propagated between recordings or exported as a CSV file.

Scene geometry is used by location export, an analysis option that exports position, body size, and body angle in every tracked frame, along with relative distance to each region and point of interest in normalized, pixel, and calibrated units. Body length and width are calculated as four standard deviations of the mask along and across its long axis.

#### Requirements and installation

Detail is supported on macOS, Linux, and Windows. It requires Python 3.12 or higher and is installed with pip or uv. Model backends and the GUI are optional extras, so a user can install only what they need. SAM 3 requires an NVIDIA GPU and CUDA.

All other models (DINOv3 and YOLO) can be run on Apple Silicon GPUs, CUDA, or a central processing unit (CPU; not recommended). Step-by-step installation instructions are in the documentation (detail-embedding.github.io/detail).

Detail guides the user through accessing SAM 3 and DINOv3 model weights, which are available through Hugging Face and require a Hugging Face account, an access token, and approved access to each model.

### Data pipeline

Here, we describe the relevant model details, parameters, and algorithms used in Detail’s data pipeline. All parameters and models described are the default, recommended settings used to generate the results presented, unless otherwise specified. All DINOv3 models in this paper are ViT-L/16 with 16-pixel patches [13]. For exhaustive documentation, see our documentation page (detail-embedding.github.io/detail).

#### Video preprocessing

At import, a timestamp for every frame is read from the video file’s presentation timestamps and stored as int64 nanoseconds. Timestamps are used as a reference across the data processing and analysis pipeline to keep every output aligned to the original video frames. Detail supports continuous recordings split across multiple files and recordings with variable frame rates.

#### Object detection

Detection is run on every video frame by default. If batch inference is supported by the detection model, frames are batched. Batch inference is not supported for SAM 3. We save the bounding box, mask, and confidence score for all detections above the confidence threshold. The confidence threshold default is 0.5. Detail currently supports detection with SAM 3 [12] using text prompts or Ultralytics YOLO segmentation models [30], either trained within Detail or imported into the tool.

#### Object tracking

The tracking stage is designed to remove spurious detections and improve the overall quality of masks. We currently support single-object tracking. Our default tracking method is the anchor tracker, a custom tracking algorithm designed to take advantage of duplicate and partial detections to produce a single object mask. The anchor tracker works as follows for each frame:

1. If a field of view exists, remove all detections whose bounding box center is outside the field of view.
2. Select the subject as the highest-confidence detection at or above the minimum confidence threshold (0.1). If no detections meet the threshold, the frame has no subject.
3. Merge up to two secondary detections, in order of confidence, with a confidence at or above the secondary confidence threshold (0.05) and an intersection over union (IoU) with the subject of at least 0.1. Leave all remaining detections unassigned.

Each frame is processed independently. Subject IoU is not calculated between frames. We also provide a simple single-subject IoU tracker.

#### Keypoint detection

Keypoint detection requires a completed detection run. Keypoints can be detected using our patch matching method or with a YOLO keypoint detection model, either distilled within Detail or imported. Detail comes with four common skeletons, and users can easily edit, create, or import their own keypoint skeletons using the Detail GUI. Keypoint detection via patch matching uses DINOv3 patch embeddings to detect keypoints across a video. The process is as follows:

1. Users mark the location of keypoints on a small number of frames (one is often sufficient).
2. DINOv3 generates patch embeddings for a cropped image of each detected subject in every frame. The crop is a square around the detection’s bounding box, resized to 448 pixels.
3. For each keypoint, we compute the cosine similarity between each patch embedding inside the subject’s mask and the embedding of the user-marked keypoint in each example frame, keeping each patch’s highest similarity.
4. The five most similar locations for each keypoint are kept as candidates and refined to sub-patch precision.
5. Final keypoint locations are chosen jointly from the candidates using the skeleton: distances between connected keypoints are kept close to those in the example frames.

For keypoint detection with a YOLO model, the model is given the entire video frame and we save every detected pose with a confidence above the confidence threshold (0.25).

#### Keypoint tracking

Keypoint tracking is designed to remove incorrect key-point detections and smooth keypoint jitter. Keypoint tracking requires a completed object tracking and key-point detection run.

The first step in keypoint tracking is assignment. Each keypoint detection is scored by the proportion of its keypoints that fall inside a subject’s mask from object tracking, after the mask is expanded by 5%. If at least half of the keypoints are within the mask, the detection is assigned to that subject. Detections not assigned to any subject are discarded.

After assignment, the keypoint trajectories are smoothed first with a Hampel filter [36] and then with a Gaussian-weighted quadratic filter. The Hampel filter replaces a point with its windowed median (*±*0.1 s) if it is more than three scaled median absolute deviations from the median, with a minimum deviation of 3% of body size. Finally, points are replaced by a quadratic curve fitted to temporally neighboring frames, with each frame weighted by a Gaussian of its time distance (*σ* = 0.133 s).

#### Embedding

Embedding is the final step in the data processing pipeline, where we create a pose image for every frame in which the subject is tracked and generate a pose embedding for that image using DINOv3. Embedding requires a completed object tracking run.

The pose image is created as follows:

1. Overlay the masked pixels of the subject on a square image canvas.
2. Set the canvas to be the color most different (maximum distance in CIELAB [37] color space) from the subject’s average color in the recording.
3. Egocentrically align the mask using a keypoint tracking run (recommended but optional).
4. Center the mask on the canvas at its true scale. The canvas is sized to the largest mask in the recording, so all masks within a recording keep their relative scale.
5. Resize all pose images to the model input size (224 pixels).

We then use DINOv3 to generate embeddings of all pose images, saving the model’s CLS token as a single 1,024-dimensional pose embedding per pose image.

#### Object detection model distillation

Model distillation is composed of two phases: dataset creation and model training. During dataset creation, the user provides a detection text string and SAM 3 generates detections for 1,000 frames by default, sampled across the selected recordings in proportion to their length and spread evenly in time within each video. Detections with a confidence above 0.5 are saved as polygon labels. Frames where SAM 3 detects nothing are kept as background examples, up to 10% of the dataset. For validation, 15% of the labeled frames are held out. Users can also create a training dataset from the saved masks of an existing detection or tracking run.

During model training, a pretrained YOLO segmentation model (YOLO11s-seg) is fine-tuned on the labeled dataset. The default training parameters are 100 epochs with a batch size of 32 and image size of 640 pixels. Detail saves the weights from the epoch with the highest box mAP50 on the validation set.

#### Keypoint detection model distillation

Keypoint model distillation follows the same two phases as object detection. The patch matching model labels keypoints on 1,000 frames by default, using the user-annotated examples (~1–5). Frames are sampled across the selected recordings in proportion to video duration and the number of frames with a tracked object. For validation, 15% of the labeled frames are held out. Users can also create a training dataset from an existing keypoint tracking run.

A pretrained YOLO pose model (YOLO11s-pose) is fine-tuned on the labeled dataset. The default training parameters are 100 epochs with a batch size of 32 and image size of 640 pixels. Detail saves the weights from the epoch with the highest keypoint mAP50 on the validation set.

#### The pose embedding space

To better understand what information embeddings capture in addition to pose (Table 4), we compared embeddings of masked and aligned pose images, masked but unaligned pose images, and full video frames from the 50 fentanyl and naloxone recordings, sampling 2,000 evenly spaced frames per recording.

We trained an MLP with one hidden layer (512 units) to predict position (mask centroid) and orientation (direction from the tail base and midpoint key-points to the head keypoint) from each embedding type, with weight decay tuned on training rats only. One female and one male rat were held out per fold over five pairings, and we report *R*^2^ on held-out rats. To test animal identity, an MLP was trained to classify the 10 rats on four treatment sessions, tested on the fifth, and scored by balanced accuracy. Own-recording share is the fraction of a frame’s 10 nearest neighbors by cosine similarity, within the same treatment and excluding frames from the same recording within 30 s, that come from its own recording. Confidence intervals are 95% intervals from 1,000 bootstrap resamples of animals.

### Analysis modules

Detail provides two analysis modules that use the pose embedding space to enable unsupervised and supervised behavior discovery and labeling. Temporal clustering produces hierarchical clusters of behaviors. Behavior querying takes user-supplied example behaviors to find similar behaviors across an experiment.

#### Temporal clustering

The objective of temporal clustering is to segment videos into short bouts of discrete behaviors and then cluster similar behaviors. A bout is a short, continuous sequence of pose embeddings corresponding to sequential frames in a video. To create behavioral bouts, videos are first split on missing embedding frames (a frame that could not be embedded because the subject was not tracked). Next, Detail computes 700 “prototype behaviors”, distinct embeddings that represent unique behaviors in the pose embedding space. To generate prototype behaviors, we compute a smoothed, windowed average embedding (0.53 s window) for every frame. Because the smoothed embeddings are computed using a windowed average across nearby frames, they contain temporal as well as pose information. We then evenly sample all smoothed embeddings, run k-means clustering [38] with *k* = 700, and average the smoothed embeddings within each group to generate our 700 prototype behavior embeddings.

Once we have produced our behavior prototypes, Detail labels every frame of an experiment with a prototype. To do this, we fit a hidden Markov model (HMM) [39] to the experiment using the smoothed embeddings. Next, we use the Viterbi algorithm [40] to find the best-scoring sequence of labels under the HMM. Each sequence of consecutive frames with the same label is a bout, and bouts shorter than 0.27 s are discarded.

Once every frame is labeled, the number of clusters is reduced via average-linkage clustering to build the hierarchical clusters that allow users to discover and label high-level behaviors such as “rearing” and sub-behaviors like “rearing on wall” or “unsupported rearing”. We use a bout’s corresponding data, including experiment metadata, video frames, and segmen-tation masks, to provide a rich suite of interpretation tools for analyzing and labeling the results of clustering.

#### Behavior querying

The objective of behavior querying is to find instances of a specific behavior across an experiment. A user selects one or more example clips of the behavior, and the pose embeddings of those clips are used to search all selected recordings for similar behavior. Detail treats querying as a frame classification problem by computing a score for every frame, computing matches based on periods of high similarity, and presenting matches ordered by confidence.

Similar to temporal clustering, the embedding sequences are first split on gaps in tracking so that no match spans a period where the subject was not tracked. Detail takes a sample of 500 “background” frames uniformly from the searched recordings, excluding the example clips. Each frame is then scored with a soft k-nearest-neighbor vote [41]. A frame’s 20 nearest neighbors among the example and back-ground frames are found by cosine similarity. Each neighbor is weighted by a softmax of its similarity (temperature 0.07), and the frame’s score is the total weight of its example-frame neighbors. This produces a score between zero and one for every frame. We smooth scores with a moving average whose length is proportional to the median length of the example clips.

Matches are based on the peaks of the smoothed score. Starting from the highest peak, each match is grown outward while the score stays above half of its peak value, up to a maximum of four times the median example length. Nearby weaker peaks are suppressed so that matches do not overlap. Each match is assigned its peak score as its confidence.

All matches are provided to the user, who can choose the confidence cutoff for each behavior after the search, reviewing the matches that fall above and below it. Similar to temporal clustering, we provide interpretation tools that use experiment metadata to support reviewing and selecting the appropriate confidence cutoff to label a behavior.

#### Auditing

Temporal clustering and querying support auditing. Results are audited by reviewing every clip of a labeled behavior (every clip of a labeled cluster in temporal clustering and every match above the chosen confidence cutoff in behavior querying). Clips the user removes count as incorrect and the rest as correct, giv-ing an exact precision. In temporal clustering, a label’s precision is the average of its clusters’ precisions, weighted by their number of clips.

### Comparisons

Comparisons measure each behavior’s share of time, rate (bouts per second), total time, and mean bout duration in each recording, averaged within each animal by default. Detail selects the matching design from the user’s grouping variable and choice of observation. Variables that are the same for all of an animal’s recordings (e.g. sex) are compared between animals by the log ratio of group means. Variables that change within an animal are compared within each animal, by the mean log ratio for two levels or the mean Spearman correlation for a numeric variable (e.g. dose). Within-animal effects are ranked by a moderated t-statistic [42], so behaviors that change consistently across animals rank highest.

*P* values come from permutation tests that shuffle group labels between animals, flip the sign of each animal’s effect, or shuffle each animal’s levels. Family-wise error is controlled by comparing each behavior’s statistic to the largest statistic across all behaviors in each permutation (200 by default, or every sign flip for 10 or fewer animals). Movement and body shape measures from location exports are each tested on their own with at least 10,000 permutations, and a Benjamini–Hochberg false discovery rate [43] is reported across measures.

### Data export

Detail supports comprehensive, granular exporting for all data created in the tool. Users can export an entire experiment, and every piece of data created during the processing pipeline and analyses. Exports include all figures, images, and plots, as well as the data used to create the visuals and the parameters of the run that produced them.

Masks and detections are exported in COCO format [44], tracks as CSV, and pose embeddings as NumPy arrays [45]. Keypoint detections can be exported as CSV or in DeepLabCut [1], SLEAP [2], or Neurodata Without Borders (NWB) [46] formats. Analysis results can be exported as video clips of each behavior bout, ethograms, and per-frame behavior labels with times-tamps, ready to be aligned with physiological or other time series data.

Users can also export an entire experiment as a single archive for backup or to be imported back into Detail by collaborators. Videos can be included in a full-experiment export but are not by default.

### Escalating fentanyl injections and naloxone-precipitated withdrawal

#### Animals

For all experiments, male and female Long-Evans rats (Charles River) weighing 250–450 g were used. Rats were dual-housed in a temperature- and humidity-controlled room on a 14:10 h light:dark cycle and acclimated to the facility for 1 week before experimental procedures. Food and water were available ad libitum. Experimental procedures were approved by the University of Washington Institutional Animal Care and Use Committee.

#### Fentanyl injections

We recorded 10 rats (five female, five male) from above for 30 min, 5 min after a subcutaneous injection of saline, fentanyl citrate (25, 50, or 75 *µ*g/kg), or naloxone (1 mg/kg). Injections were given twice daily, with the recording after the second (afternoon) injection. On the naloxone day, a 75 *µ*g/kg fentanyl dose was given in the morning, followed by the naloxone dose for the afternoon injection and recording. Every rat was tested in every condition.

#### Recordings

Rats were placed individually in a gray plastic bin (floor, 45.7 *×* 30.5 cm) and filmed from above for 30 min at 1,920 *×* 1,080 pixels and 62.5 frames per second with a global shutter camera (Arducam B0610).

#### Segmentation and tracking

Videos were processed with Detail, and locomotion and body shape were measured from Detail’s tracked masks with a custom Python script. A segmentation model (YOLO11s) distilled from SAM 3 detections from these recordings segmented the rat in every frame, and a keypoint detection model (YOLO11s-pose) located the head, the middle of the body, and the base of the tail. The rat was tracked in at least 99.1% of each session (median 99.8%). For the measures below, the tail was removed from each mask (morphological opening with a disc whose diameter was 12% of the square root of that animal’s median mask area), and masks were sampled five times per second.

#### Locomotion

Position was the center of the mask, smoothed with a 1 s running average to remove frame-to-frame jitter. Distance traveled is the summed length of the path. The path was broken, not joined, across gaps in tracking and across apparent jumps of more than 1.5 body lengths between samples. A rat was counted as moving when its speed exceeded 2 cm/s. Time moving is the percentage of tracked time above that speed, and speed while moving is the mean speed over that time. Occupancy maps show the time spent in each 1.2-cm bin, averaged across rats.

#### Body shape

Body length and width are four standard deviations of the mask’s pixels along and across its long axis, and the length-to-width ratio was computed in each frame. Each measure was summarized as its median over the session.

#### Behavior clusters

Behaviors were identified without labels by temporal clustering with default settings, pooling all 50 recordings. Clusters with at least 30 s of total time were screened for differences between conditions with the comparisons module, and clusters that differed were named by an experimenter after viewing their most typical bouts.

#### Statistics

All comparisons were within animal (*n* = 10) and used two-sided permutation tests, which make no assumption about the distribution of the data. For locomotion and body shape, the effect of dose was tested with the Spearman correlation between dose (saline counted as 0) and the measure; the reference distribution was built by shuffling each rat’s four sessions 20,000 times. Saline and naloxone were compared with a paired sign-flip test on the mean within-rat difference. For behavior clusters, the share of session time was compared with Detail’s comparisons module, as a trend over dose and as a paired contrast of nalox-one versus saline, with family-wise correction across clusters (200 permutations for dose and all 1,024 sign flips for naloxone). Clusters seen in fewer than half of the rats were not tested, leaving 89 clusters for dose and 105 for naloxone.

## Discussion

Pose estimation is the predominant method for quantifying animal behavior, and provides accurate, interpretable measurements, but comes with some practical limitations. Pose estimation models require substantial human annotation to achieve acceptable accuracy in each experimental setting, and the small number of keypoints commonly used to represent pose may constrain our ability to capture the continuum of animal behavior. Detail, in its present form, is not a replacement for pose estimation. Its keypoints, placed by DINOv3 patch matching from a few annotated frames, are designed to orient each animal for egocentric alignment before embedding, so that clustering in the embedding space reflects posture rather than position and orientation. The rise of foundation vision models presents an alternative pathway for automated behavior quantification and analysis, giving scientists the ability to generate rich, detailed representations of pose for any animal, in any setting, with minimal or no annotation.

Detail is our attempt to provide a foundational platform to access and research embedding-space-derived behavioral analyses. While several existing methods have explored using foundation vision models and image embeddings to capture animal behavior, Detail brings the entire workflow together into a single, unified environment. Within Detail, video frames are mapped into a high-dimensional embedding space that captures rich, detailed, and scientifically useful encodings of pose. To harness these embeddings, Detail provides a user-friendly interface, an efficient and species-agnostic workflow, and integrated analytical modules. This allows behavioral scientists to easily process hundreds of hours of footage and annotate, discover, and explore behavior all in one place.

We have engineered our data pipeline to be model-agnostic, allowing the accuracy of segmentation, keypoint detection, and image embedding to improve at the rapid pace of frontier model development. Detail was designed for flexibility and built to evolve as the field matures, offering a central, adaptable resource, and a complete alternative or complement to keypoint-based analytical methods.

Detail’s highly generalizable data pipeline is designed for a wide range of behavioral analysis applications. The primary limitations of our current data pipeline are video quality and multi-animal experiments. Detail can only represent the behavioral dynamics that are captured in experimental footage. While segmentation models such as SAM3 are highly robust to varying image quality, bad camera angles, blurry or obscured footage, and messy video scenes still interfere with the ability of segmentation models to reliably isolate subjects and reduce the quality of behavioral information that can be represented in pose embeddings. Multi-animal support is a high priority and under active development. Lastly, pose embeddings are less directly interpretable than keypoints, because measures such as joint angles and distances cannot be read from them, and (common to all un-supervised approaches) behaviors found by clustering must be reviewed and named by a person. Current development of Detail is focused on introducing features to help automate and standardize the descriptions of Detail’s unsupervised behavioral clusters.

With the rise of larger, more computationally demanding machine learning models, computational efficiency is an increasingly relevant issue for automated behavior analysis. Detail is designed to allow users with a wide range of resources to benefit from the full capabilities of the tool. Detail can be run effectively on a single workstation with an NVIDIA GPU. Using smaller YOLO models produced by our built-in model distillation pipeline, Detail can be run at slower speeds on even smaller machines like modern Mac-Books. Detail can also take full advantage of more powerful multi-GPU machines, and support for cloud-based compute resources and HPC clusters is under active development.

To facilitate ease of use and adoption, we provide extensive documentation at detail-embedding.github.io/detail. Our code is available at github.com/detail-embedding/detail. Detail is an ongoing project, and we aim to build a research community around Detail to support the development of new capabilities, improve accuracy and efficiency, and expand its use to new domains. We welcome bug reports, feature requests, and contributions through the GitHub repository.

## Acknowledgments

We are grateful to Zhihan Zhang, Sewoong Oh, Jason Ren, and Adam Pikielny for their helpful discussions about embedding spaces.

## Declarations

### Consent for publication

All authors consent to publication.

### Data availability

The data used in this study are available at github.com/detail-embedding/detail.

### Code availability

Detail is available at github.com/detail-embedding/detail, with documentation at detail-embedding.github.io/detail.

### Author contributions

Conceptualization: M.L., K.R.C., S.A.G. Data curation: K.R.C., S.A.G. Formal analysis: M.L., K.R.C., S.A.G. Funding acquisition: V.I., K.R.C., S.A.G. Investigation: M.L., K.R.C., S.A.G. Methodology: M.L., K.R.C., S.A.G. Project administration: K.R.C., S.A.G. Resources: V.I., R.J., K.R.C., S.A.G. Software: M.L., K.R.C., S.A.G. Supervision: V.I., R.J., K.R.C., S.A.G. Validation: M.L., K.R.C., S.A.G. Visualization: M.L., K.R.C., S.A.G. Writing – original draft: M.L. Writing – review & editing: M.L., V.I., R.J., K.R.C., S.A.G.

